# Sympatric versus allopatric evolutionary contexts shape differential immune response in *Biomphalaria / Schistosoma* interaction

**DOI:** 10.1101/378034

**Authors:** Anaïs Portet, Silvain Pinaud, Cristian Chaparro, Richard Galinier, Nolwenn M. Dheilly, Julien Portela, Guillaume M. Charriere, Jean-François Allienne, David Duval, Benjamin Gourbal

**Affiliations:** Univ. Perpignan Via Domitia, Interactions Hôtes Pathogènes Environnements UMR 5244, CNRS, IFREMER, Univ. Montpellier, F-66860 Perpignan, France; Interactions Hôtes-Pathogènes-Environnements (IHPE), UMR 5244, CNRS, Ifremer, Université de Perpignan Via Domitia, Université de Montpellier, Montpellier, 34095, France.; School of Marine and Atmospheric Sciences, Stony Brook University, Stony Brook, New York, USA.

## Abstract

Selective pressures between hosts and their parasites can result in reciprocal evolution or adaptation of specific life history traits. Local adaptation of resident hosts and parasites should lead to increase parasite infectivity/virulence (higher compatibility) when infecting hosts from the same location (in sympatry) than from a foreign location (in allopatry). Analysis of geographic variations in compatibility phenotypes is the most common proxy used to infer local adaptation. However, in some cases, allopatric host-parasite systems demonstrate similar or greater compatibility than in sympatry. In such cases, the potential for local adaptation remains unclear. Here, we study the interaction between *Schistosoma* and its vector snail *Biomphalaria* in which such discrepancy in local versus foreign compatibility phenotype has been reported. Herein, we aim at bridging this gap of knowledge by comparing life history traits (immune cellular response, host mortality, and parasite growth) and molecular responses in highly compatible sympatric and allopatric *Schistosoma/Biomphalaria* interactions originating from different geographic localities (Brazil, Venezuela and Burundi). We found that despite displaying similar prevalence phenotypes, sympatric schistosomes triggered a rapid immune suppression (dual-RNAseq analyses) in the snails within 24h post infection, whereas infection by allopatric schistosomes (regardless of the species) was associated with immune cell proliferation and triggered a non-specific generalized immune response after 96h. We observed that, sympatric schistosomes grow more rapidly. Finally, we identify miRNAs differentially expressed by *Schistosoma mansoni* that target host immune genes and could be responsible for hijacking the host immune response during the sympatric interaction. We show that despite having similar prevalence phenotypes, sympatric and allopatric snail-*Schistosoma* interactions displayed strong differences in their immunobiological molecular dialogue. Understanding the mechanisms allowing parasites to adapt rapidly and efficiently to new hosts is critical to control disease emergence and risks of Schistosomiasis outbreaks.

**Author summary:** Schistosomiasis, the second most widespread human parasitic disease after malaria, is caused by helminth parasites of the genus *Schistosoma*. More than 200 million people in 74 countries suffer from the pathological, and societal consequences of this disease. To complete its life cycle, the parasite requires an intermediate host, a freshwater snail of the genus *Biomphalaria* for its transmission. Given the limited options for treating *Schistosoma mansoni* infections in humans, much research has focused on developing methods to control transmission by its intermediate snail host. *Biomphalaria glabrata*. Comparative studies have shown that infection of the snail triggers complex cellular and humoral immune responses resulting in significant variations in parasite infectivity and snail susceptibility, known as the so-called polymorphism of compatibility. However, studies have mostly focused on characterizing the immunobiological mechanisms in sympatric interactions. Herein we used a combination of molecular and phenotypic approaches to compare the effect of infection in various sympatric and allopatric evolutionary contexts, allowing us to better understand the mechanisms of host-parasite local adaptation. Learning more about the immunobiological interactions between *B*. *glabrata* and *S*. *mansoni* could have important socioeconomic and public health impacts by changing the way we attempt to eradicate parasitic diseases and prevent or control schistosomiasis in the field.

## Introduction

Schistosomiasis is the second most widespread human parasitic disease after malaria and affects over 200 million people worldwide [1]. *Schistosoma mansoni* (Platyhelminthes, Lophotrochozoa) causes intestinal schistosomiasis. *Schistosoma* needs a fresh water snail acting as its first intermediate host to undergo part of its life cycle before infecting humans. Patently infected snails support the continuous production of thousands of cercariae, infective for humans. Vector snails are central actors of the parasite transmission and obvious targets for schistosomiasis control that deserve more attention. It is therefore necessary to understand snail-parasite immunobiological interactions and to characterize the molecular mechanisms of successful snails and *Schistosoma* interactions.

The compatibility of numerous strains of *Biomphalaria glabrata* and *Schistosoma sp.* has been extensively tested, revealing that (i) different *B. glabrata* laboratory strains (or isolates) show various degrees of susceptibility to *S. mansoni* infection and (ii) different strains of *S. mansoni* display different levels of infectivity towards a particular strain of snail host [2–6]. Compatibility is defined as the ability for the miracidia to infect snail and become a living primary sporocyst in snail tissue. Incompatibility refers to miracidia that are recognized by the snail immune system and encapsulated and killed by the hemocytes (the snail immune cells). Thus, the success or failure of the infection of *B. glabrata* by *S. mansoni* reflects a complex interplay between the host’s defense mechanisms and the parasite’s infective strategies, based on a complex phenotype-to-phenotype or matching-phenotype model [2–4, 7–9]. In the past 15 years, the molecular basis of this compatibility polymorphism has been investigated at the genomic [10–12], transcriptomic [8, 13–17], proteomic/biochemical [18–23] and epigenomic levels [24–29]. These studies have revealed that various molecules and pathways involved in immune recognition (snail immune receptors versus parasite antigens), immune effector/anti-effector systems, and immune regulation/activation participate in a complex interplay that governs the match or mismatch of host and parasite phenotypes [30]. This complex phenotype-by-phenotype interaction or compatibility polymorphism varies between populations and individuals resulting in a “multi-parasite susceptibility” or “multi-host infectivity” phenotypes [4] that reflect between-population variations in parasite infectivity/virulence and host defense/resistance [31, 32].

Most of the time, interaction in *B.glabrata/Schistosoma* models has been investigated by comparing, (i) sympatric/compatible and (ii) allopatric/incompatible host-parasite associations. The general assumption is that the parasites thanks to their shorter generation times, larger population sizes and higher reproductive outputs, are ahead in the co-evolutionary race against their host and are therefore more likely to locally adapt and perform better when infecting local hosts [33, 34], than allopatric hosts [34–37]. However, in many instances, Schistosomes are highly compatible to hosts from other localities, showing the same or even greater infection success when exposed to allopatric hosts. Thus they do not fulfil the “local versus foreign” main criterion of the local adaptation between a host and its parasite [5, 38–40]. Very few studies have investigated the molecular basis of allopatric compatible interactions from the perspective of both side of the interaction, the host and the parasite [41, 42].

Hence, in order to bridge this gap, we herein study sympatric/allopatric interactions displaying similar compatibilities using an integrative approach that links the underlying molecular mechanisms to the resulting phenotypes, based on comparative molecular approaches on both host snails and *Schistosoma* parasites. We characterize the underlying cellular and molecular mechanisms of the interaction between South American snail strains (from Recife Brazil and Guacara Venezuela) and three different highly compatible parasite isolates: (i) the sympatric strains of *S. mansoni* from Recife Brazil, (ii) the allopatric *S. mansoni* from Guacara Venezuela (narrow geographic scale), and (iii) the allopatric *S. rodhaini* from Burundi Africa (large geographic and phylogenetic scales).

Our results clearly show that even though the compatibility phenotypes among these strains is similar, a very different immunobiological dialogue is taking place between *B. glabrata* vector snails and their sympatric or allopatric *Schistosoma* parasites at the cellular and molecular levels.

## Results

### A RNAseq approach of host immune response in sympatric and allopatric infections

The *B. glabrata* transcriptome was analyzed using the previously described RNAseq pipeline developed in our laboratory [8, 43, 44]. Of the 159,711 transcripts of the *Bg*BRE transcriptome, 3,865 (2.4%) were differentially represented in all sympatric and allopatric conditions compared to naive snails (Table 1, S1 Fig). We performed automatic Blast2GO annotation, discarded the non-annotated transcripts, and retained 1,017 annotated transcripts (26.3% of the differentially expressed (DE) transcripts, S1 Fig). In the following analysis, we focused on the 336 transcripts known to have immune-related functions (8.7% of DE transcripts, S1 Fig).

**Table 1:** Number of transcripts in each step of transcriptomic analysis

| Analysis of transcriptomic data |  | Analyses of Blast2GO | Analyses of annotations |  |
| --- | --- | --- | --- | --- |
| Full Transcriptome | Differentially expressed | Informative annotation | Immune Transcripts | Non Immune Transcripts |
| 159,711 | 3,865 | 1,017 | 336 | 681 |

Of these immune related transcripts, 189, 180, and 164 DE transcripts were identified in the BB (BgBRE/SmBRE, sympatric), BV (BgBRE/SmVEN, allopatric), and BR (BgBRE/Srod, allopatric) interactions, respectively (Fig 1A). Among those, 40 transcripts were consistently differentially expressed in response to infection (Fig 1A). They also displayed similar expression profile (Fig 1B, cluster 1).

**Fig 1:**
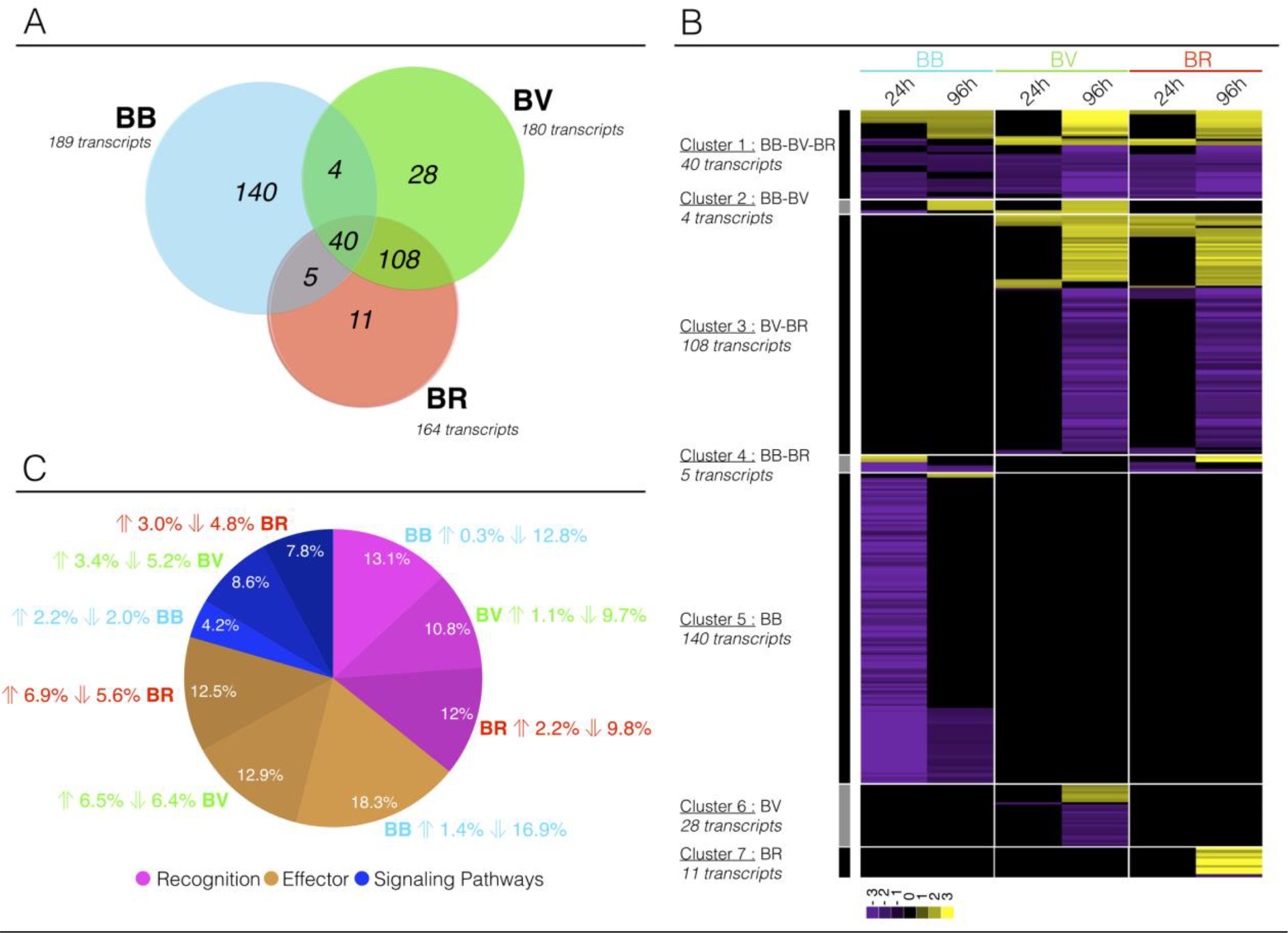
Dual-RNAseq of Biomphalaria immune-related transcripts. Among the differentially represented transcripts, Blast2GO functional annotation allowed us to identify 336 transcripts that appeared to be related to the *Biomphalaria* immune response. Abbreviations and colors: blue BB, sympatric interaction between BgBRE and SmBRE; green BV, allopatric interaction between BgBRE and SmVEN; and red BR, allopatric interaction between BgBRE and Srod. For each interaction 40 whole-snails are used, 20 pooled at 24h and 20 at 96h post-infection. A) Venn diagram showing the relationships among the immune transcripts found to be differentially expressed in the sympatric and allopatric interactions. B) Clustering of differentially represented immune transcripts. Heatmap representing the profiles of the 336 differentially represented immune-related transcripts in the BB, BV, or BR interactions along the kinetic of infection (at 24 and 96 h). Each transcript is represented once and each line represents one transcript. Colors: yellow, over-represented transcripts; purple, under-represented transcripts; and black, unchanged relative to levels in control naïve snails. C) Pie chart showing the distribution of the selected immune-related transcripts across three immunological processes: immune recognition (pink), immune effectors (brown), and immune signaling (blue). For each category and interaction, the respective proportion of transcripts and the direction of the effect (over- or underexpression) are indicated.

Most (74.1%) of the transcripts differentially expressed in response to infection by the sympatric parasite (BB) were not differentially expressed in response to either one of the two other parasites. Most importantly, all of the sympatric-specific transcripts were under-represented at 24 h post-infection, and 74.6% of these transcripts were differentially expressed exclusively at this time point (Fig 1B, cluster 5), suggesting a parasite-induced immunosuppression.

In contrast, very similar transcript expression patterns were observed in response to infection by the two different species of allopatric parasites: *S. mansoni* (BV) and *S. rodhaini* (BR) and most of the variations in gene expression occurred 96h after infection. Of the 108 transcripts consistently differentially expressed in allopatric response (Fig 1A; Fig 1B, cluster 3), 98.1% were differentially expressed at 96 h post-infection, and 28.2% were more abundant following infection (Fig 1B, cluster 3). Transcripts differentially expressed exclusively in response to SmVEN or Srod were groupd in Clusters 6 (28 transcripts) and 7 (11 transcripts), respectively. In response to SmVEN (BV, Fig 1B, cluster 6), 96.5% of the transcripts were differentially abundant 96 h after infection (22% over-represented) and in response to Srod (BR, Fig 1B, cluster 7), 100% of the transcripts were differentially abundant 96 h after infection (82% over-represented).

We explored the function of DE transcripts in response to the three different parasites. We initially distributed the relevant differentially expressed immune transcripts into three groups: (i) immune recognition molecules, (ii) immune effectors, and (iii) immune signalling molecules (Fig 1C, S2 Table), that were then subdivided into functional categories (Fig 2). When we compared the percentage of each immunological group in the sympatric and allopatric interactions, no specific functional subset was particularly repressed in the BB sympatric interaction (Fig 1C; Fig 2). The same immune functions were affected in response to infections by sympatric or allopatric parasites but different immune transcripts (grey and black diamond in Fig 2) showed differential regulation following infections (Fig 2).

**Fig 2:**
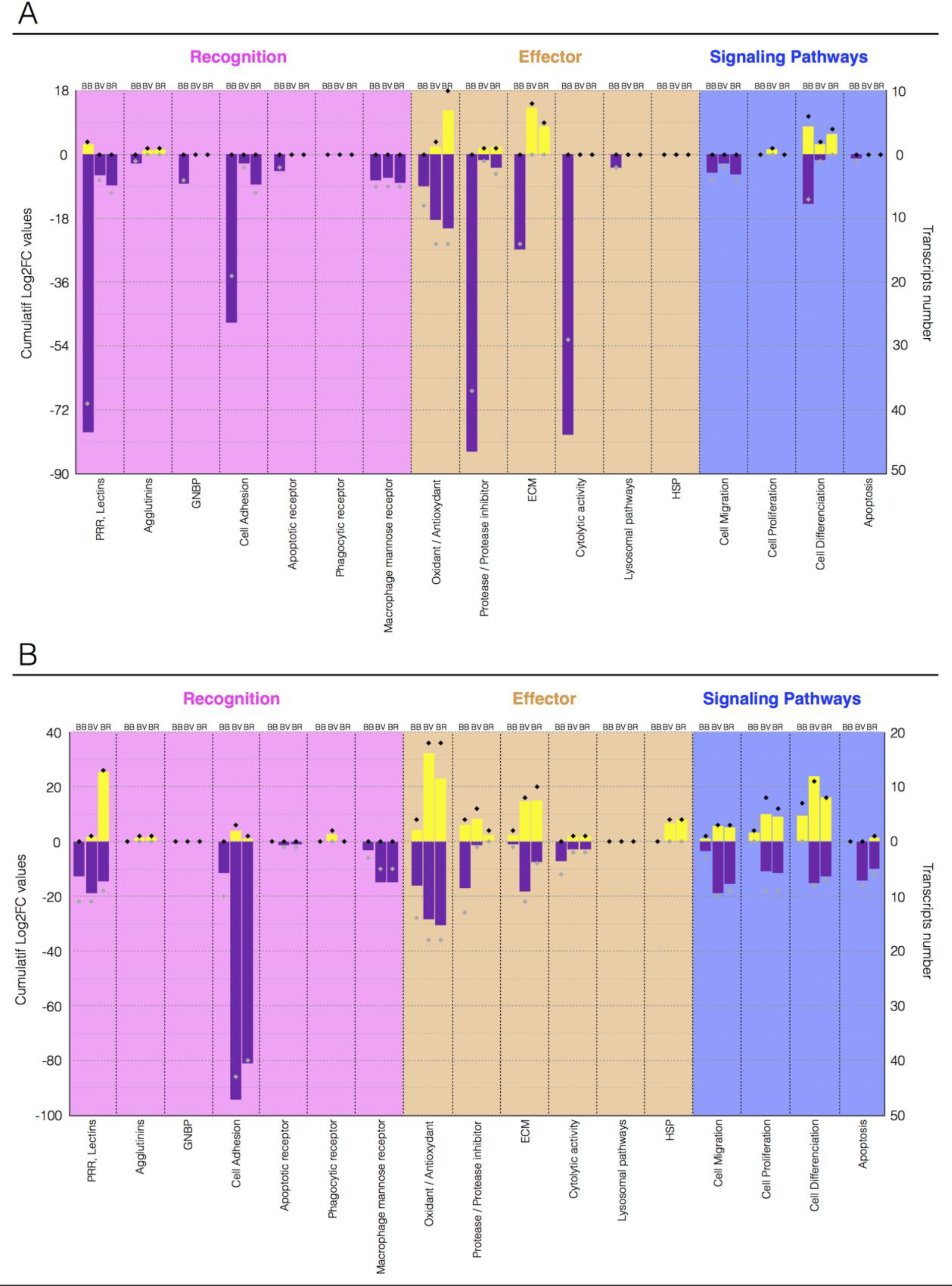
Differentially represented immune-related transcripts in sympatric and allopatric interactions. Cumulative expression [Log2FC (fold change) from DESeq2 analysis] of the immune-related transcripts identified as being differentially represented following sympatric or allopatric infection. Transcripts were grouped into the three immunological groups described in Fig 1, and from there into functional categories. The yellow histograms correspond to cumulatively over-represented transcripts, while the purple histograms show under-represented transcripts. The black (over-represented) and gray (under-represented) diamonds correspond to the number of transcripts analyzed in each functional category. Abbreviations: BB, BgBRE/SmBRE interaction; BV, BgBRE/SmVEN interaction; and BR, BgBRE/Srod interaction. A. Immune transcript expression at 24 h post-infection. B. Immune transcript expression at 96 h post-infection.

The differentially regulated transcripts belonging to the three immunological groups (Fig 2) were largely involved in immune cellular responses, cell adhesion, extra cellular matrix component, cell migration, cell differentiation and cell proliferation. These functions were consistently reduced at the 24h time point in sympatric interaction (76%), whereas many transcripts involved in the same molecular processes were over-represented in allopatric interactions (39%) (Fig 2).

### Immune cellular responses in the sympatric and allopatric contexts

Hemocytes, the snail immune cells, participate directly in the immune response against the parasites, and immune cell activation under an immunological challenge can translate into cell proliferation and/or cell morphology modifications. Thus, cell proliferation was quantified using *in vitro* (Fig 3) and *in vivo* (Fig 4) EdU nuclear labelling. EdU is a nucleoside analogue of thymine incorporated into DNA during DNA synthesis. Its incorporation reflects the mitotic activity of hemocytes.

**Fig 3:**
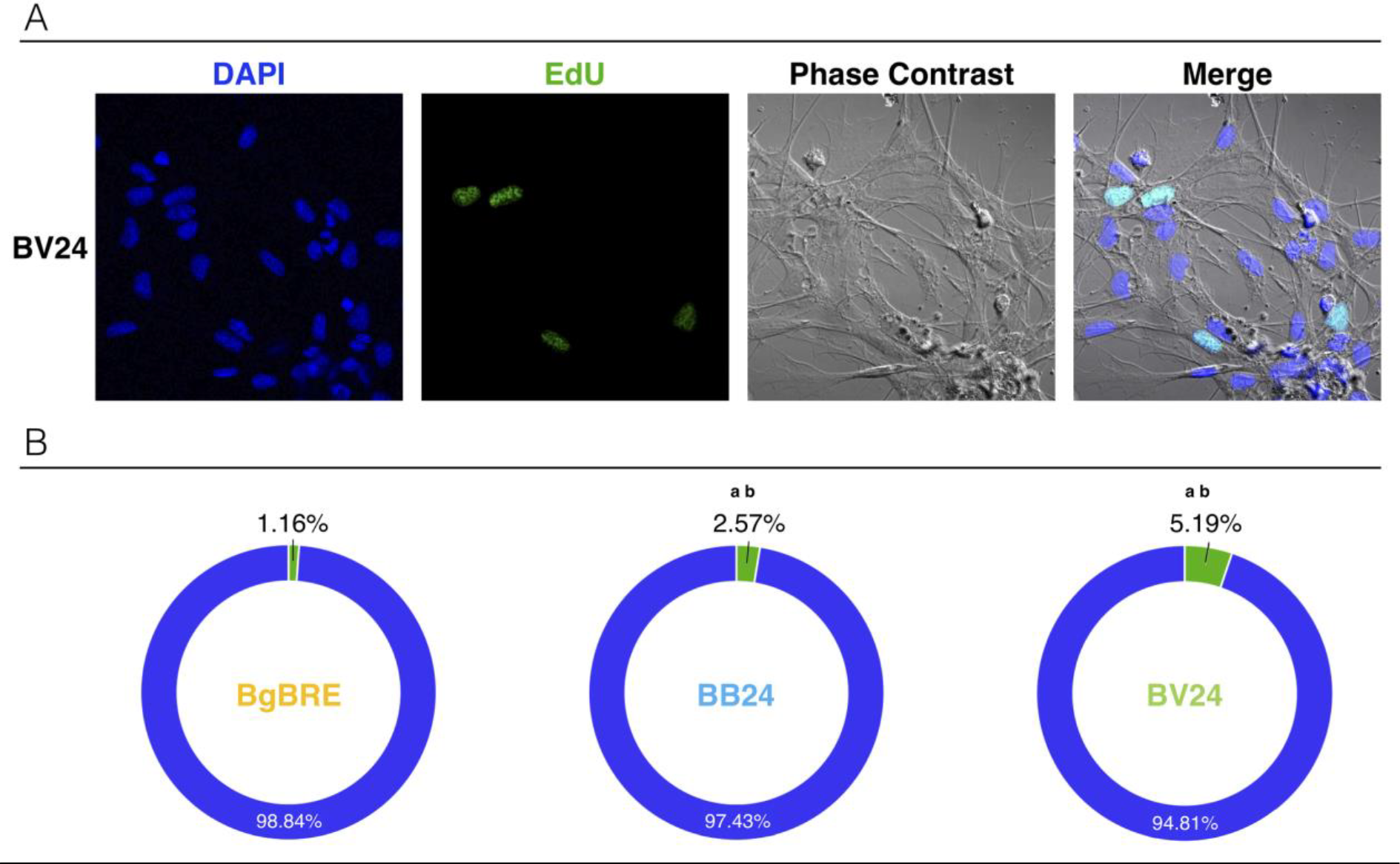
Microscopic analyse of snail hemocyte proliferations. In vitro EdU labeling of hemocytes was conducted for sympatric and allopatric interactions A) Hemocytes were collected at 24 h post-infection for *in vitro* analysis. Confocal microscopy of EdU-labeled hemocytes from snails subjected to the allopatric BV interaction (BgBRE/SmVEN). Colors: blue/DAPI; green/EdU; white/phase contrast. B) Microscopic counting of EdU-labeled hemocytes from naïve control snails (BgBRE) (n=1,811) and those subjected to the sympatric interaction (BB: BgBRE/SmBRE) (n=2,064) or an allopatric interaction (BV: BgBRE/SmVEN) (n=1,366) recovered from 3 individual snails by condition. Colors: green, EdU-positive cells; and blue, EdU-negative cells. Between-group differences in the percentage of proliferation were tested using a Fisher exact test, with statistical significance accepted at p<0.05. The “a” indicates a significant difference between the naïve and infective conditions, while “b” indicates a significant difference between the infective conditions.

**Fig 4:**
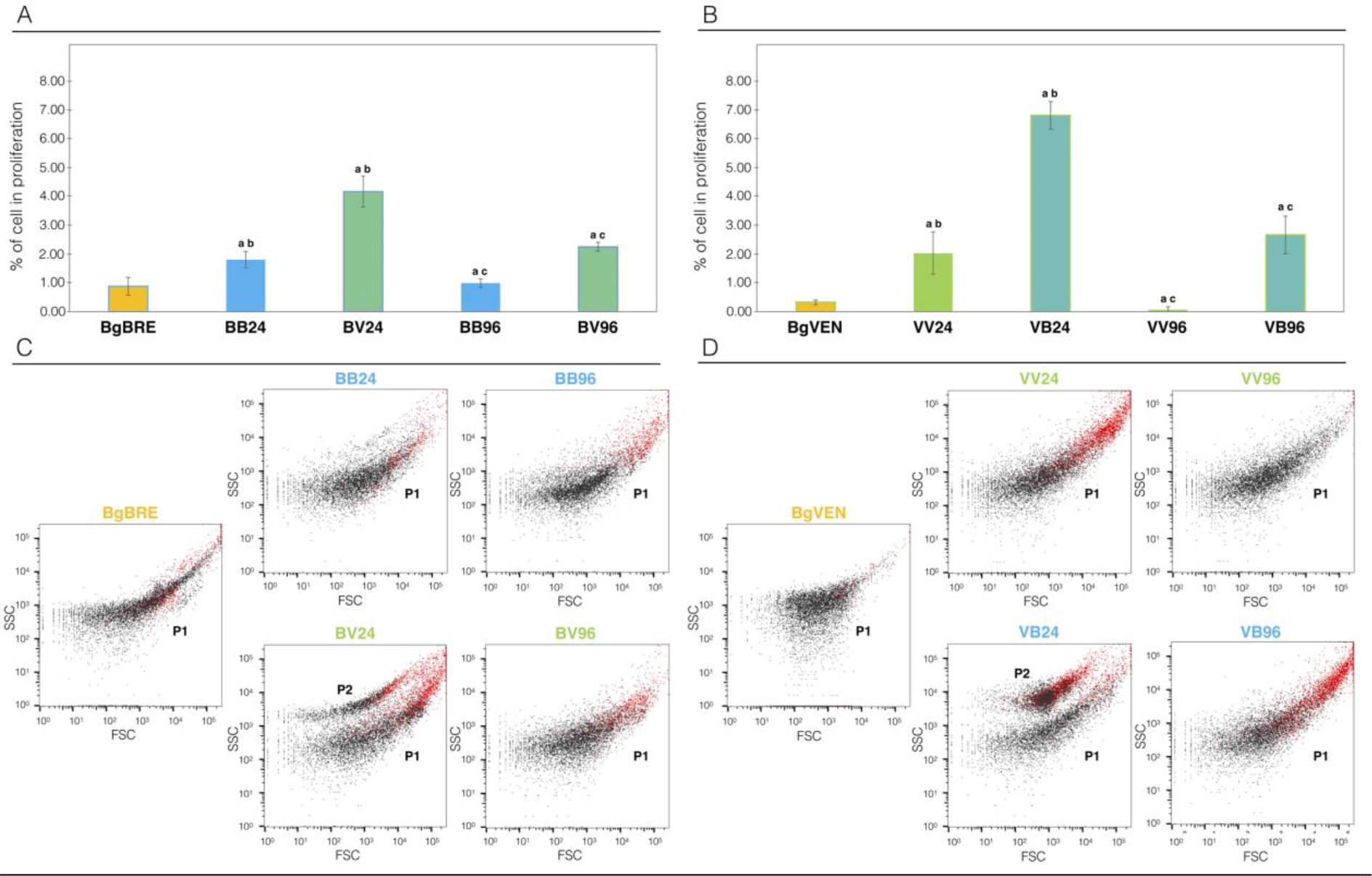
Flow cytometry analyse of the hemocyte response in sympatric and allopatric interactions. A) Flow cytometry was used to count *in vivo* EdU-labeled hemocytes at 24 h and 96 h after infection in sympatric and allopatric interactions. A total number of hemocytes of n=10,000were recoveredfor6 biological replicates of 3 snails. Control naïve snails (BgBRE, yellow) were compared to those subjected to the sympatric interaction (BB, BgBRE/SmBRE, blue) or an allopatric interaction (BV, BgBRE/SmVEN, green). B) The experiment described in A was repeated using the BgVEN snail strain. Control naïve snails (BgVEN, yellow) were compared to those subjected to the sympatric interaction (VV, BgVEN/SmVEN, green) or an allopatric interaction (VB, BgVEN/SmBRE, blue). C) FSC (forward-scattered light, representing cell size) and SSC (side-scattered light, representing cell granularity) circulating hemocyte patterns in BgBRE snails under the naïve condition (yellow) or 24 h and 96 h after infections in sympatry (BB24/96, BgBRE/SmBRE, blue) or allopatry (BV24/96, BgBRE/SmVEN, green). D) FSC and SSC circulating hemocyte patterns in BgVEN snails under the naïve condition (yellow) or 24 h and 96 h after infections in sympatry (VV24/96, BgVEN/SmVEN, blue) or allopatry (VB24/96, BgVEN/SmBRE, green). The red dots correspond to EdU-positive hemocytes. Between-group differences in the percentage of proliferation were tested using the Mann-Whitney U-test, with statistical significance accepted at p<0.05. The “a” indicates a significant difference between the naïve and infective condition, “b” indicates a significant difference between the infective conditions at 24h, and “c” indicates a significant difference between the infective conditions at 96h.

*In vitro* labelling was used on circulating hemocytes recovered from BgBRE 24h after infection with SmBRE and SmVEN to compare the proportion of mitotic circulating hemocytes in sympatric and allopatric interaction, respectively (Fig 3A). Quantification of Edu-positive hemocytes using confoncal microscopy showed that 24h after infection, hemocyte proliferation was 3 times more important following infection of BgBRE by SmVEN (5.2% of proliferative cells in BV) than SmBRE (2.6% of proliferative cells in BB) (Fisher exact test two-tailed p = 7.6 e10^−6^) (Fig 3B). Moreover, this result demonstrates for the first time that “circulating” hemocytes are able to proliferate following *Schistosoma* infections.

Hemocyte proliferation 24h after infection was then further assessed using flow cytometry after *in-vivo* EdU-labelling (Fig 4A, B).

Here, we performed the same experiments using another *Biomphalaria glabrata* strain, BgVEN as the host and SmVEN and SmBRE as the sympatric and the allopatric parasite, respectively (Fig 4B). The rate of proliferating cells was significantly higher in allopatric than sympatric interaction in both BgBRE and BgVEN (BgBRE Mann Whitney U test: U=36; z = −2.8; p = 0.0022; BgVEN, Mann Whitney U test: U=36; z=-2.8; p = 0.0022). In BgBRE, allopatric interaction (BV) was associated with 4.2% of proliferative cells whereas sympatric interaction resulted in 1.8% of proliferative cells (Fig 4A). In BgVEN, allopatric interaction (BgVEN/SmBRE, VB) was associated with 6.8% of proliferative cells whereas sympatric interaction (BgVEN/SmVEN, VV) resulted in 2.0% of proliferative cells (Fig 4B). At 96 h after infection, there were fewer proliferating cells: the percentage of proliferating hemocytes in sympatric BB and VV interactions were similar to the non-infected controls (BB, 1%, Mann Whitney U test: U=17; z=-0.27; p = 0.3936; VV, 0.1%, Mann Whitney U test: U=2; z=2.48; p = 0.013), while remaining somewhat higher in both allopatric interactions (BV, 2.3%, Mann Whitney U test: U=0; z=2.65; p = 0.009; .VB, 2.7%, Mann Whitney U test: U=36; z=2.8; p = 0.0022). These results confirm that the reduced cell proliferation is associated with sympatric interaction regardless of the strain used.. The morphology of hemocytes (size and granularity) from non-infected and infected *B. glabrata* snails (BgBRE and BgVEN) in sympatric and allopatric interactions with the parasites SmBRE and SmVEN was observed using flow cytometry (Fig 4C and 4D). Morphology and heterogeneity of circulating hemocytes varied similarly in BgBRE and BgVEN snails (Fig 4C and 4D). In non-infected snails, the content of circulating hemocytes was very heterogeneous, but represented a single population with continuous gradient of size and granularity typical of *B. glabrata* hemocytes (Fig 4C and 4D) [45]. Hemocyte population heterogeneity changed quickly after infection. In allopatric interactions, 24 h after infection (Fig 4C, BV24, and 4D, VB24) two populations could be distinguished: a population P1 (corresponding to that seen in non-infected snails) and a population P2 (a new population). P2 cells exhibited increased granularity, retained a high degree of size variability, and showed a mitotic activity, as indicated by EdU labeling (Fig 4C and 4D, red dots). This profile was transitory, as the P2 population had disappeared 96 h after infection (Fig 4C, BV96, and 4D, VB96). Altogether, these results show that, upon infection, the snail circulating immune cells exhibit a particular population dynamic with transient increase of the mitotic activity associated with morphology modifications. Moreover, this cellular response appears to be inhibited by sympatric parasites.

### *Schistosoma* growth and development in *Biomphalaria* tissues

#### Parasite development

To investigate the development of *S. mansoni* in *B. glabrata* tissues, we examined the fate of sporocyst in sympatric and allopatric compatible interactions using a histological approach, for this we used 3 snails per conditions. For both interactions, miracidia were able to penetrate, transform into primary sporocysts (SpI), and develop. At 24 h after infection, we observed a significant difference (Mann Whitney U test: U=40; z=4.33; p = 1.42 e10^−6^) in the size of sporocyst from sympatric parasites (11,838 µm^2^ average size on 9 parasites) versus allopatric parasites (7,402 µm^2^ average size on 8 parasites) (Fig 5). A small difference in sporocyst size is still observed at 96 h after infection but without being significant (41,413 µm^2^ on 7 parasites for sympatric and 36,920 µm^2^ on 10 parasites for allopatric, Mann Whitney U test: U=280; z=-1.31; p = 0.1917) (Fig 5). These results show that during the early events following infection, the allopatric parasites develop more slowly than sympatric one’s; thereafter, allopatric parasites seemed to catch up quickly, resulting in no significant difference in size observed at 96 h post-infection (Fig 5).

**Fig 5:**
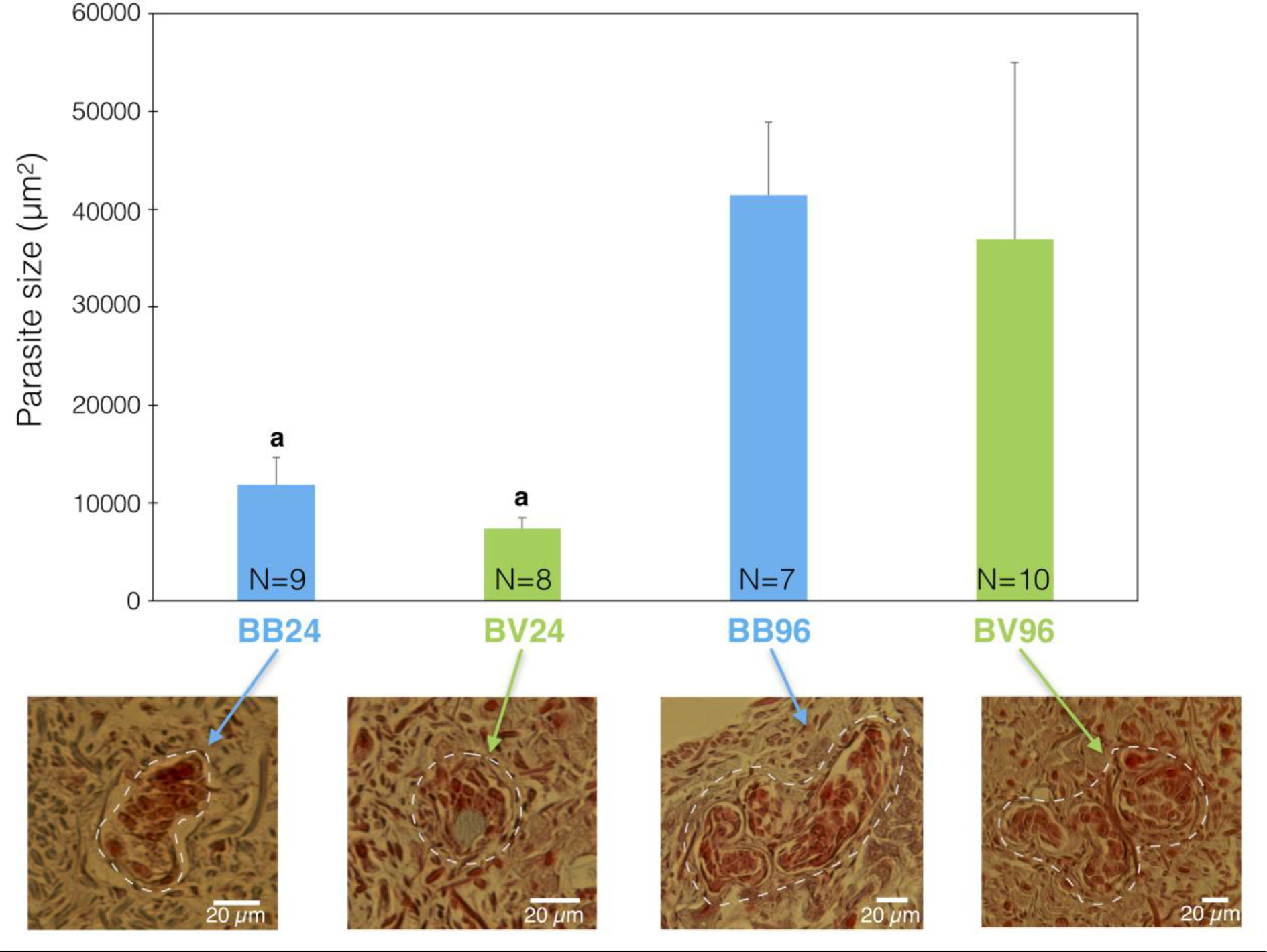
Development of parasites into snail tissues. A histological approach was used to monitor parasite size along the course of snail infection. The sympatric interaction (BgBRE x SmBRE) is shown in blue, and the allopatric interaction (BgBRE x SmVEN) is shown in green. For each experimental interaction, the parasite sizes were quantified at 24 and 96 h after infection. Morpho-anatomical aspects of the parasite are depicted to highlight a potential difference in parasite survival. N=7 to 10sporocystes were used as indicated in the figure.Between-group parasite size differences were assessed using the Mann-Whitney U-test, with significance accepted at p<0.05 (indicated by “a” on the histograms).

#### Parasite transcript expression analysis

We used dual RNAseq data to identify transcripts expressed by SmBRE, SmVEN and Srod during their intra-molluscal development in BgBRE. The parasite RNAseq data at 24 h after infection, revealed five clusters of DE transcripts from the sympatric (SmBRE) and the allopatric (SmVEN, Srod) parasite responses (Fig 6). Cluster 1 corresponds to transcripts highly expressed and cluster 5 weakly expressed for all parasite strains. Cluster 2 represents transcripts over-expressed in SmBRE versus SmVEN and Srod. Cluster 3 contained transcripts over-expressed in SmBRE and SmVEN versus Srod and cluster 4 SmBRE and Srod versus SmVEN. In all clusters, the transcript expression levels in SmBRE sympatric parasite are always greater than for the other allopatric parasites. Blast2GO annotation was successful for 70% of the 351 transcripts identified in the five clusters described above (S3 Table). According to the global Gene Ontology (GO): 70% of the annotated genes were involved in general metabolism and growth, translation processes, regulation of cellular processes and RNA biosynthesis; 25% were involved in molecular transport or cell organization; and 5% were involved in organism defence or response to stimuli. In all these clusters, we identified 6 parasite gene products been involved in parasite modulation or suppression of snail immunity. These molecules correspond to heat shock proteins (Fig 6, clusters 1 and 2) [27]; glutathione-S-transferase, NADH dehydrogenase subunit, and calreticulin (Fig 6, cluster 2) [20, 46, 47]; Alpha-2-macroglobulin (Fig 6, cluster 4) [48]; von willebrand factor type EGF with pentraxin domain (Fig 6, cluster 5) [49] (see S3 Table).

**Fig 6:**
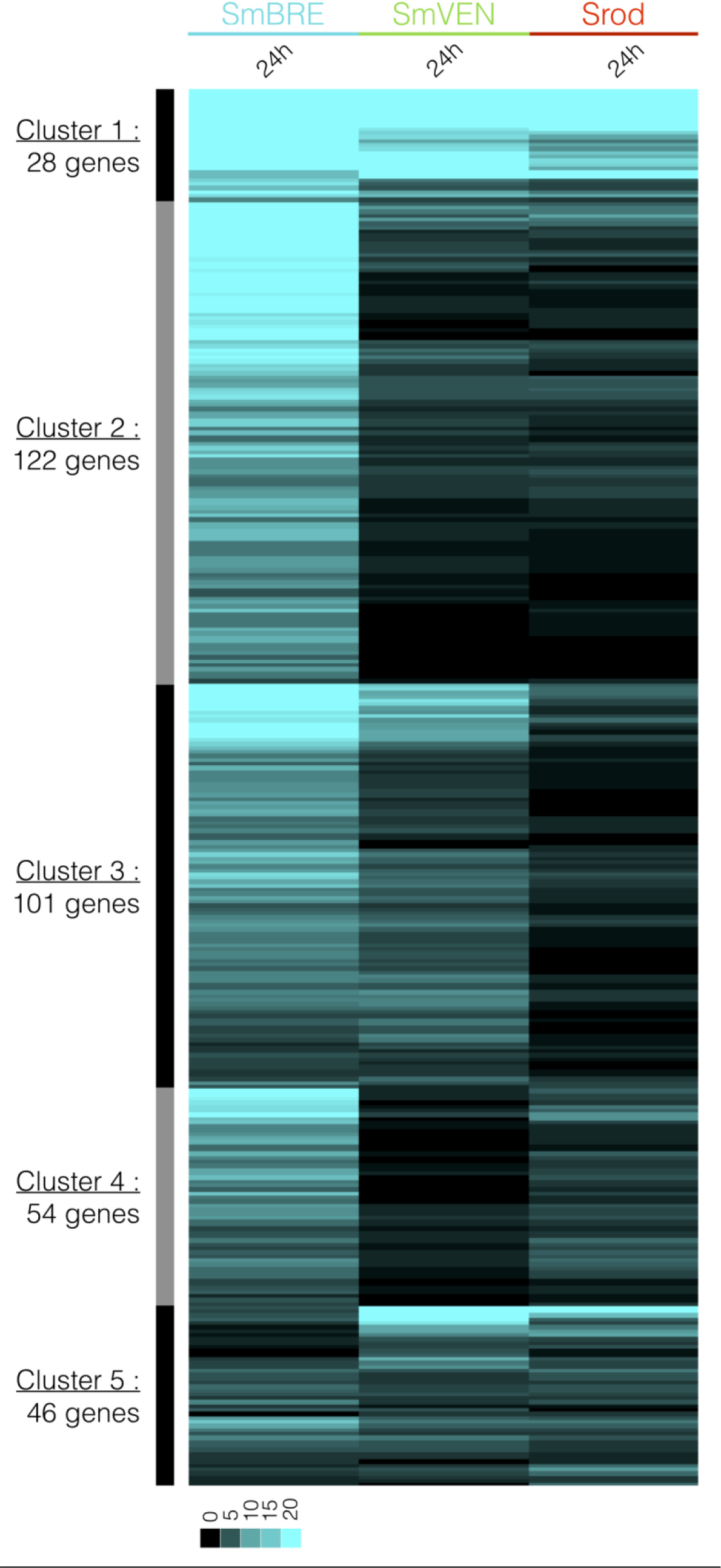
Clustering of intra-molluscalSchistosoma expression patterns. RNAseq library mapping enabled us to identify 351 genes expressed by *Schistosoma* parasites in *Biomphalaria* snail tissues. Colors: blue, *S. mansoni* Brazil (SmBRE); green, *S. mansoni* Venezuela (SmVEN); and red, *S. rodhaini* (Srod). The heatmap represents the profiles of the 351 genes expressed by the different parasites at 24 h after infection. Each transcript is represented once and each line represents one transcript. The expression level is highlighted by the different shades of blue.

Interestingly, allopatric parasites did not over express any transcripts that could have immunosuppression function or impair the activation of the immune response (Fig 6 and S3 Table). Furthermore, a variant of a glycerol-3-phosphate acyl-transferase (Schisto_mansoni.Chr_3.5623) is highly over expressed in SmVEN and Srod compared to SmBRE (cluster 1, S3 Table). This molecule is known to participate in the biosynthesis of phosphatidic acid, itself involved in macrophage activation and regulation of inflammatory signalling [50, 51].

#### Parasite microRNAs analysis

The microRNAs (miRNAs) are known as non-coding small RNA (<24nt) highlighted to regulate gene expressions. As we identified strong differences in the transcriptional responses between sympatric and allopatric interactions, we can hypothesized that the processes of transcriptional or post-transcriptional regulations may be deeply affected. In this respect, we investigated *in-silico,* the potential presence of *Schistosoma mansoni* miRNAs (sma-mir) in our parasite RNAseq data. At 24 h post-infections, we identified 54 miRNA precursors from miRBase with high quality alignment scores against the different RNAseq read libraries (naïve BgBRE, BB24, BV24, BR24). To avoid cross-species misidentifications, we selected precursors that were exclusively identified in infected and never identified in uninfected snails (naive BgBRE). Eleven miRNA precursors corresponding to *Schistosoma mansoni* were identified (Fig 7A). Nine of the parasite miRNA precursors were specific to the Brazil-infected libraries (BB24); two were specific of the Venezuela-infected libraries (BV24); and one was shared across the three infected conditions (BB24, BV24 and BR24). Although we identify 49 miRNA precursor sequences specific to *S. mansoni* (Fig 7B), we decided to select only miRNAs covered by 100% nucleotide similarity that allowed to predict 11 miRNAs in mature (eg. sma-mir-2d-3p, sma-mir-190-3p) or precursor (sma-mir-8431) forms. Then, in order to identify candidate sequences that could represent putative miRNA targets, we used the Miranda tool (S4 Table). Only RNA-RNA interactions that showed good scores for pairing (>140) and enthalpy (<15 Kcal) were considered. The number of targets pertaining to the differentially expressed immune-related transcripts identified in Fig 1 that were found for the identified miRNAs ranged from 2 targets for sma-mir-8456, to 50 targets for sma-mir-2d of the differentially expressed immune-related transcripts.

**Fig 7:**
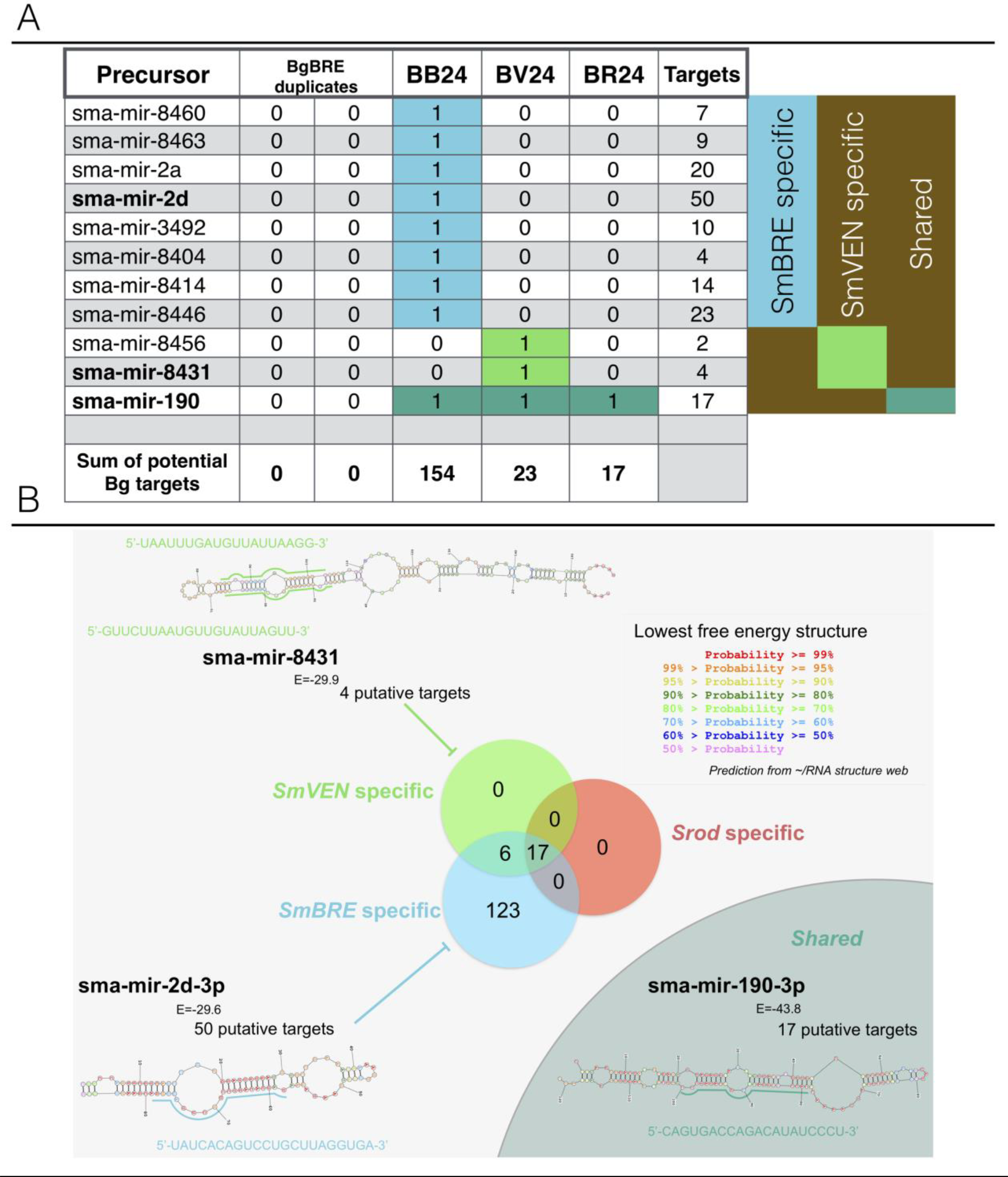
In-silico identification of parasite miRNAs. miRNAs were assessed using libraries obtained from naïve snails and snails infected for 24 h under the various interaction conditions (BB24, BgBRE x SmBRE; BV24, BgBRE x SmVEN; BR24, BgBRE x Srod). A) Table highlighting the precursor miRNAs that may have targets among the immune-related snail transcripts selected in the present work. They include eight precursors specifically recovered in BB24, two in BV24, and one shared across the three infected conditions. The total numbers of potential targets in each condition are indicated. B) Venn diagram showing the potential targets according to the sympatric or allopatric interactions. Shown is an example miRNA stem-loop precursor that presents the highest number of potential host targets.

The miRNAs identified under the sympatric condition (SmBRE) were predicted to potentially target 43.5% of the differentially represented immune-related transcripts identified in the RNAseq experiment (Fig 1B, Fig 7) whereas 6.8% and 5.1% were targeted in allopatric conditions, SmVEN and Srod, respectively with fewer available miRNA as well (Fig 1B, Fig 7). The lack of such potential weapon to target host immune system in allopatric compared to the sympatric strain may explain the absence of immunosuppression observed in allopatric conditions. Otherwise, we did a focus on miRNAs that were shared between sympatric and allopatric interactions to try to understand the similar prevalence observed between sympatric and allopatric infections. Like so, we identified one miRNA: sma-miR-190-3p (Fig 7 and S4 Table). This miRNA was predicted to bind 17 different targets among which, we identified different variants of the Fibrinogen Related Protein (FREP) family and a cytotoxic/cytolytic humoral factor the biomphalysin. To go further, we look at the expression of those candidates following infection. If FREP transcripts were down regulated in sympatric interaction, it is not always the case in allopatry. However, interestingly all biomphalysin transcripts were under-represented in sympatric and allopatric interactions. Altogether, these data suggest that the parasites might hijack the host immune response using dedicated miRNAs as the sma-miR-190-3p described in the present study.

### Survival of snail following infection

To examine the potential impact of allopatric or sympatric parasites on snail survival, we investigated the mortality rates of infected snails over 4 months. The survival rate was significantly higher for non-infected snails compared to infected snails (sympatric interaction Kaplan-Meier Log Rank test p = 1.39 e10^−5^ and allopatric interaction p = 0.0005). However, there was no significant difference in the mortality rates of snails subjected to sympatric versus allopatric interactions: at the end of the experiment, the survival rates were 72% and 65% for the allopatric and sympatric interactions, respectively (Kaplan-Meier Log Rank test p = 0.243) (S2 Fig).

## Discussion

In the natural environment, it is assumed that the parasitic genes responsible for infectivity will evolve alongside the host defence genes, resulting in adaptation of the interactions between local host and parasite populations [52, 53]. In this context, local/sympatric parasites were expected to display a greater infectiveness, reproductive success, and virulence in host populations compared to foreign/allopatric parasites [36, 37, 54, 55]. However, in some cases this rule may be contradicted, as certain allopatric parasite-host interactions have been reported to be significantly more compatibles than the corresponding sympatric combinations [56, 57], it appears that certain *Biomphalaria/Schistosoma* interactions do not fulfil at the local adaptation between the host and the parasite, in which the sympatric parasite is expected to perform better than the allopatric one [36, 37, 54, 55].

Using field data, Morand et al. (1996) [38], Prugnolle et al. (2006) [5] and Mutuku et al. (2014) [39] showed that although sympatric parasite-host combinations of schistosomes and snails do tend to be more compatible, exceptions exist wherein particular allopatric combinations are equally or significantly more compatibles. Similar results were obtained when comparing the interactions of Brazilian and Guadeloupean snails versus *Schistosoma* infections [41]. The authors found that allopatric Guadeloupean parasites were not able to infect Brazilian snails; but Brazilian parasites were able to infect the allopatric Guadeloupean snails. Furthermore, this work demonstrated the presence of local adaptation between reactive oxygen species (ROS) and ROS scavengers in this system [41]. Based on these observations, we propose that it would be important to develop integrative analysis to depict and understand the precise molecular crosstalk (immunobiological interactions) occurring in such highly compatible sympatric and allopatric systems. Thus, dual-comparative approaches were used herein to simultaneously analyze the responses of *Biomphalaria* snails and *Schistosoma* parasites into sympatric or allopatric interactions displaying similar compatibilities.

The present RNAseq analysis demonstrated that in sympatric interaction (BB) a huge immunosuppression occurs. Twenty-four hours after the infection, the three immunological processes: (i) immune recognition, (ii) effector and (iii) signaling pathways (Fig 1 and 2) were down regulated. Conversely, in allopatric interactions (BV and BR), host immune response was activated after 96 hours (Fig 1 and 2). Differentially regulated transcripts mostly belong to immune cellular activation, migration, proliferation, or differentiation (Fig 2). An EdU labelling was used to detect proliferation and confirmed that hemocyte proliferation is inhibited during interaction with two different strains from Brazil and Venezuela (Fig 3, 4A and 4B). In addition, we discovered that a new subpopulation of proliferating hemocytes (named P2), exclusively differentiate 24h following allopatric infections (Fig 4C and 4D). P2 was EdU-positive and characterized by an increased in granularity, indicating that the new P2 cell subtype could proliferate (Fig 4). However, in absence of specific hemocyte markers, it is difficult to analyze precisely which hemocyte morphotype are proliferating (Fig 4C, D). The P2 subpopulation would thus originates from either a morphological change in an existing subset (correlating potentially with a decline in the P1 population), or represents cells that are migrating from tissues or hematopoietic organ to reach the hemolymph. Indeed, P2 population reflects newly proliferating cells that present higher EdU positive cells than the P1 population (Fig 4C, D). Further investigations will be necessary to conclude on the origin of P2 population.

In *Biomphalaria* snails, we know 3 main hemocyte morphotypes, the blast-like cells, the type I hyalinocytes and the granulocytes [58]. Based on the flow cytometry and Edu labelling approaches we can supposed that bigger and granular cells (granulocytes and hyalinocytes) are the ones that proliferates. This is demonstrated in S3 Fig in which Edu labelling was observed for hyalinocytes and granulocytes but never for blast-like cells (S3 Fig). These results seem to demonstrate a differentiation or sub-functionalization in hemocyte subtypes following infection.

This differentiation or sub-functionalization is different comparing sympatric and allopatric interactions, i.e., hemocyte proliferation decreased more rapidly in sympatric rather than in allopatric interactions (Fig 3 and 4), P2 population observed solely in allopatric interactions (Fig 4). Using reciprocal sympatric and allopatric interactions, we demonstrate that the cellular or molecular phenotype observed refers to potential co-evolution or adaptation rather to a simple host or parasite strain effect (Fig 3 and 4).

The strong immunosuppression observed within 24h of infection by a sympatric parasite, and the inhibition of hemocyte proliferation can certainly explain the differences in the growth of sympatric and allopatric parasites. Indeed, we observed a significant difference in sporocyst size 24h after infection (Fig 5), with sympatric sporocysts that were one-third bigger than allopatric sporocysts. But, 96h after infection, there was no more significant size difference between sympatric and allopatric parasites (Fig 5). This difference in size between the sympatric and the allopatric parasites at the beginning of the interaction can be explained by several hypotheses, (i) a delay in development of the allopatric parasite due to the necessity to circumvent the host immune response, (ii) the intrinsic ontogenesis or morphogenesis of post-miracidial intramolluscan stages that can be longer for allopatric SmVEN parasite compared to sympatric SmBRE parasite, finally (iii) the miracidial binding and penetration into the tissues of the host may take longer for the allopatric parasite than for the sympatric parasite. The consequences of this delay in terms of secondary sporocyst development, number of cercariae produced, or cercariae infectivity and pathogenicity for the vertebrate host, will deserve further investigation to conclude about a potential fitness cost between sympatric and allopatric parasites.

To find new clues as to how sympatric parasites immunosuppress the host or circumvent the host immune system, we used a dual-RNAseq approach to compare transcripts expression of the sympatric and allopatric parasite intra-molluscal stages (Fig 6). As the histological differences were solely observed at 24h after infection, we used dual-RNAseq to investigate the parasite expression patterns at the same time point of infection. Most of the parasite transcripts belonged to the processes of nucleotide metabolism, transcription, translation and cell differentiation, development, and growth. We also identified some transcripts with GO terms or functions related to immuno-modulation or immuno-suppression (Fig 6 and S4 Table). Nearly all of the identified transcripts were over-represented in the sympatric interaction compared to the allopatric ones. Our results therefore suggest that the installation, development and growth of the parasite occurred much more rapidly in the BgBRE/SmBRE combination, as sympatric parasites seemed to interfere more efficiently with the host immune system. However, RNAseq data did not give any clear information about how allopatric parasites succeed in circumventing the host immune system. We thus next examined the generated dual-RNAseq libraries in an effort to identify whether sympatric and/or allopatric schistosomes could hijack the host immune system using microRNAs. To begin testing this hypothesis, we confronted the dual-RNAseq data to the *Schistosoma mansoni* subset of miRBase to identify the presence of parasite microRNAs (pmiRNAs) in our datasets. Even if we don’t know whether pmiRNAs were present in contact with the host immune system or simply endogenic, this in-silico exploration may ask the question to a potential molecular discussion between metazoan organisms in a host-parasite system, based on nucleic acid weapons. miRNAs are known to regulate numerous biological processes, including key immune response genes [59, 60]. Recent work has demonstrated that circulating small non-coding RNAs from parasites have hijack roles against host metabolism, notably in the interaction of schistosomes with their vertebrate hosts [61–63]. Such non-coding RNAs could act as exogenous miRNAs to interfere with or circumvent the host immune system. In the present study, 24h after infection, several differentially expressed pmiRNAs were identified. We predicted targets of such pmiRNAs in the *Biomphalaria* immune reference transcriptome and found that they may target 43.5% of the differentially regulated immune transcripts identified in the RNAseq approach (Fig 7). In contrast, far fewer correspondences were identified for the allopatric interactions (Fig 7). The higher proportion of targeted genes in the sympatric interaction may be responsible for the observed efficient immunosuppression. If confirmed, such mechanism would reveal a specific co-evolution or adaptation in the transcriptional regulation between sympatric host and parasite. However, even if more host immune genes appeared to be targeted in the sympatric combination compared to the allopatric one’s (Fig 7), both sympatric and allopatric interactions displayed the same ability to succeed to infect the host. This similarity in compatibility phenotype between sympatric and allopatric parasites could potentially results from their ability to target host immune weapons or host genes that regulate innate cellular response using miRNAs. A unique miRNA was found in all allopatric and sympatric parasites, sma-miR-190-3p. It is predicted to bind various targets including Fibrinogen Related Protein (FREP) and biomphalysin. The FREP family members are known as pathogen recognition receptors [64, 65] and FREP knockdown is associated with an increase of snail compatibility toward *Schistosoma* infections [66, 67]. The biomphalysins belong to beta pore forming toxins and are key humoral factors of biomphalaria snails involved in cytotoxic/cytolytic activities against *Schistosoma* parasites with the ability to bind miracidia and sporocyst surfaces [68, 69]. Moreover, transcription of these molecules was mostly reduced in sympatric and allopatric interactions (figs. 1 and 2) supporting the hypothesis that sma-miR-190-3p or other pmiRNA members could play an essential role in parasite compatibility. Parasites expressing such miRNAs would thus be considered as highly virulent parasites with strong infecting capabilities. By producing dedicated miRNAs, the parasites were potentially able to regulate transcriptional, post-transcriptional, translational and protein stability processes that might help them to subvert the snail’s immune defences. Even if these results are particularly interesting, a dedicated small RNAs (<30nt) sequencing is now mandatory to validate or not the miRNA molecular cross talk occurring between Schistosome larval stages and their snail intermediate hosts as it has been shown for the interaction with their vertebrate definitive hosts.

Compatibility reflects the outcome of complex immunobiological interactions and depends on: (i) the ability of the snail immune system to recognize and kill the parasite; and (ii) the ability of the parasite to circumvent or evade the host immune response [20, 46, 70]. Based on the present observations, we propose that sympatric and allopatric interactions trigger totally different responses. In the sympatric interaction, the parasite is able to induce a host immunosuppression within the first day of infection enabling it to quickly infect the host and readily begins its development. In the allopatric interaction, the parasite is not able to quickly neutralize the host immune system, and as a consequence the parasite is recognized by host defense system that mounts a potent immune response. In allopatric parasite, the disruption of the activation of their developmental program during the first day of infection could results from the need to resist to the snail immune system. However, they seemed to be able to quickly protect themselves against the host immune response and develop normally in snail tissues as soon as 96h post-infection. Thereafter, in the medium- or long-term, there are no observable differences in the prevalence, intensity, or snail survival comparing sympatric and allopatric interactions (S1 Table, S2 Fig).

Thus, we show that despite having similar prevalence phenotypes, sympatric and allopatric snail- *Schistosoma* interactions displayed a very different immunobiological dialogue at the molecular level. Intriguingly, these different immunobiological interactions seem to have no repercussions upon parasite growth at longer term or to host survival. As differences at the molecular level do not correspond apparently to any ecologically meaningful changes in term of fitness, it is not straightforward to demonstrate local adaptation in such systems. However, we do not know if fitness costs could affect other biological traits in sympatric and allopatric interactions, as for example secondary sporocysts production and growth, number of cercariae shedding, or cercariae infectivity and pathogenicity towards the vertebrate host. Demonstrating local adaptation would thus appear extremely complex and would indeed deserve further investigation. It is hard to draw the line as to when local adaptation is or is not present. However, our results argue that the differences find at the molecular level may ultimately contribute to the evolution of local adaptation at an ecological level.

Nevertheless, the ability for allopatric pathogens to adapt rapidly and efficiently to new hosts could have critical consequences on disease emergence and risk of schistosomiasis outbreaks.

Past events of allopatric parasites reaching new areas of transmission, even in large-geographic scale dispersion, have been largely documented. The most famous example being the schistosomiasis colonization of South America since the slave trade of the 16^th^-19^th^ Centuries [71, 72]. *Schistosoma* originated in Asia, reached Africa 12 to 19 million years ago (MYA), and gave rise to all *Schistosoma* species known in Africa [72]. *S. mansoni* diverged from *S. rodhaini* around 2.8MYA [71, 73], and thereafter,400 to 500 years ago, colonized South America [71, 72]. This colonization of South America by *S. mansoni* from Africa was rendered possible by the presence of the snail host: *Biomphalaria glabrata*. All African species of *Biomphalaria* are monophyletic and seem to have originated from paraphyletic South American clade [74–76]. The ancestor of *B. glabrata* appears to have colonized Africa 1 to 5 MYA, giving rise to all 12 species of *Biomphalaria* known today in Africa [77]. In South America and Caribbean Island, *S. mansoni* infects *B. glabrata*; in Africa, it infects mostly *B. pfeifferi* and *B. alexandrina*. We found that South American *S. mansoni* parasites are highly compatible with their sympatric South American snail hosts, whereas African *S. mansoni* parasites display low compatibility phenotype with South American snail hosts (S1 Table). Interestingly, the South American parasites did not lose their compatibility for African snail hosts; i.e., the prevalences are similar to African parasites when confronted to African snails (S1 Table). The recent African origin of South American *Schistosoma* parasites (introduction in South America 400 to 500 years ago) may explain why they have not diverged sufficiently in South America to lose their compatibility for African snail hosts. In this case, the transfer of allopatric parasites from Africa to South American snail hosts have be successful and result in the emergence of schistosomiasis in South America.

More recently another case of compatible allopatric parasite emergence have been observed when schistosomiasis have reach Europe [78, 79]. Here, humans infected in Senegal have imported a hybrid between *Schistosoma haematobium* and *Schistosoma bovis* into Corsica. In this case urogenital schistosomiasis could be introduced and easily and rapidly spread into this novel area of south Corsica because *Bulinus truncatus* the vector snail of *S. haematobium* was endemic in the Corsica Cavu River [78, 79]. However, this allopatric African hybrid parasite was able to adapt efficiently to the Corsican new *B. truncatus* host. If parasite hybridization can potentially have a putative role in increasing the colonization potential of such *S. haematobium*, it would be particularly interesting to analyze and depict the molecular support of such allopatric interactions to predict the potential risk of schistosomiasis outbreaks in other European areas, or other potential transmission foci.

If we hope to draw conclusions regarding the existence of emerging or outbreak risks, we need to develop integrative approaches to explore fine-scale patterns of host-parasite interactions. We must consider the spatial scale at which comparisons are conducted, the patterns of disease occurrence, the population genetics, and the involvement of physiological, immunological, and molecular processes. Studying the relevant factors at the relevant timing would be of critical importance in terms of schistosomiasis control. Understanding further, how these allopatric parasites efficiently infect host snails would be mandatory to identify markers and develop new tools to predict or to quantify risks of schistosomiasis outbreaks. Now it would be particularly relevant to go back to the field to see how translatable are our results in a more dynamic field situations with genetically diverse hosts and parasites witch evolved under complex abiotic and biotic interactions, with newly encountered allopatric hosts and potentially on quite different spatial scales. For this we have a wonderful playground in Corsica.

## Materials and Methods

### Ethics statement

Our laboratory holds permit # A66040 for experiments on animals from both the French Ministry of Agriculture and Fisheries, and the French Ministry of National Education, Research, and Technology. The housing, breeding and animal care of the utilized animals followed the ethical requirements of our country. The researchers also possess an official certificate for animal experimentation from both French ministries (Decree # 87–848, October 19, 1987). Animal experimentation followed the guidelines of the French CNRS. The different protocols used in this study had been approved by the French veterinary agency from the DRAAF Languedoc-Roussillon (Direction Régionale de l’Alimentation, de l’Agriculture et de la Forêt), Montpellier, France (authorization # 007083).

### Biological materials

The two studied strains of *S*. *mansoni* (the Brazilian (SmBRE) or the Venezuelan (SmVEN) strains) and the strain of *S. rodhaini* (Srod) had been maintained in the laboratory using Swiss OF1 mice (Charles River Laboratories, France) as the definitive host. Two snail strains of *Biomphalaria glabrata* were used in this study: the albino Brazilian strain, (BgBRE) and the Venezuelan strain, (BgVEN). All host and parasite strains of each different geographical origin were recovered in their native locality and parasite strains were maintain in the laboratory always on their sympatric snail hosts to maintain the same selective pressure and sympatric adaptation on parasite. We housed snails in tanks filled with pond water at 25°C with a 12:12 hour light:dark cycle and supplied ad libitum with fresh lettuce. The Brazilian strain originates from the locality of Recife (east Brazil, recovered in the field in 1975), the Venezuelan strains of snail and parasite were recovered from the locality of Guacara (north Venezuela, recovered in the field in 1975) and the African species *Schistosoma rodhaini* originates from Burundi and was obtained from the British Museum National History (recovered in 1984). These *Schistosoma* isolates/species have been selected because they exhibited similar infectivity toward BgBRE or BgVEN strains (see prevalence and intensity in S1 Table). These high compatibilities were followed-up by the cercariae emissions. For all these interactions we observed comparable cercariae shedding (S1 Table). Prevalence of SmBRE and SmVEN for the African vector snail *Biomphalaria pfeifferi* from Senegal (BpSEN), and prevalence of the corresponding parasite SmSEN on South American snails were also tested (S1 Table).

### RNAseq experimental protocol

In order to investigate the molecular response of snails against sympatric and allopatric parasites, a global comparative transcriptomic approach was conducted. One hundred and twenty BgBRE snails were infected with SmBRE, SmVEN or Srod. Each snail was individually exposed for 12 h to 10 miracidia in 5mL of pond water. For each experimental infection, 30 snails were recovered at 24h and 96h after infection. Pools of 30 snails were composed of 10 juvenile snails (shell diameter from 3 to 5 mm), 10 mature adult snails (shell diameter from 7 to 9 mm) and 10 old adult snails (shell diameter from 11 to 13 mm). The samples were named as follows: BB24, BB96 for BgBRE infected with SmBRE; BV24, BV96 for BgBRE infected with SmVEN; and BR24, BR96 for BgBRE infected with Srod. We realised 2 pools of 30 uninfected BgBRE snails (pool of immature, mature and old snails) named Bre1 and Bre2, that were used as control conditions for all downstream comparative analyses.

#### Whole-snail RNA extraction and sequencing

Total RNA was extracted using TRIZOL^Ⓡ^ (Sigma Life Science, USA) according to the manufacturer’s instructions. Sequencing was performed using paired-end 72-bp read lengths on Illumina Genome Analyzer II (MGX-Montpellier GenomiX, Montpellier, France).

#### De novo transcriptome assembly

*De novo* transcriptome assembly, using all time points, was performed using an in-house pipeline created with the Velvet (1.2.02), Oases (v0.2.04), and CD-HIT-EST (v4.5.4) programs. The assembly of the consensus reference transcriptome was optimized using various parameters, including k-mer length, insert length and expected coverage, as previously described [43, 44]. A *de novo* transcriptome was created and contained 159,711 transcripts.

#### Differential expression analysis

High-quality reads (Phred score >29) were aligned to the *de novo* transcriptome using Bowtie2 (v2.0.2), which was run locally on a Galaxy server. To compare the host responses during the sympatric or allopatric interactions, we used the DESeq2 (v2.12) was used to analyse the differential transcript representation between *BgBRE* control strains (uninfected *Bg*BRE1 and *Bg*BRE2) to the sympatric and allopatric conditions (p-value < 0.1) [44]. A Venn diagram was generated using the Venny 2.1 software to highlight which differentially expressed transcripts were specific or common to the different interactions. A heatmap was obtained using the log2 Fold Change with Hierarchical Ascending Clustering (HAC) and Pearson correlation (uncentered) as applied by the Cluster (v3.0) and Java TreeView (v1.1.6r4) softwares packages. The differentially represented transcripts were functionally classified using a BlastX analysis with the cut-off set to e-value < 1e^-3^ (NCBI dataset; thanks to the Roscoff Data center Cluster, UPMC) and gene ontology was assigned using an automatic annotation, implemented in Blast2GO (v3.0.8) (S2 Table). We identified potential immune transcripts involved in snail immunity based on functional domains predictions and literature searches.

### Schistosoma intra-molluscal stage transcriptome analysis: Dual RNA-seq

A dual RNA-seq approach was conducted to gain in a broader understanding of sympatric and allopatric host/parasite interactions.

#### Schistosome read selection

The *Biomphalaria* (v1) and *Schistosoma* (v5.2) genomes were concatenated (https://www.vectorbase.org/organisms/biomphalaria-glabrata; http://www.sanger.ac.uk/resources/downloads/helminths/schistosoma-mansoni.html). Then high quality reads (Phred score >29) were mapped against these concatenated genomes using Bowtie2 (v2.0.2), run locally on the Galaxy project server. The reads that mapped only once and exclusively to the *Schistosoma* genome were collected as corresponding to *Schistosoma* reads; reads that mapped to the *Biomphalaria* genome or more than once to either genomes were removed from the analysis.

#### Gene analysis

The above-selected *Schistosoma* reads were mapped against the concatenate genome to identify intra-molluscal stage-specific *Schistosoma* genes. In order to select the relevant genes, the reads mapped in all experimental conditions were summed. Solely genes with a minimal sum of 10 reads were kept for the analysis. A heatmap was generated to analyse *Schistosoma* gene expression patterns using Hierarchical Ascending Clustering (HAC) with Pearson correlation (uncentered) as applied by the Cluster (v3.0) and Java TreeView (v1.1.6r4) software packages. Functional annotation of the genes was assigned using BlastX with the cut-off set to e-value < 1e^-3^ (NCBI dataset, local cluster) and gene ontology was performed using Blast2GO (v4.0.7) (S3 Table).

### Innate immune cellular response analysis: microscopy and flow cytometry

Hemocytes appeared as the main cells supporting *Biomphalaria* snail immune response. Thus, to go further in the description of snail response against parasites, quantitative and qualitative changes in hemocyte populations were investigated. For this purpose, BgBRE and BgVEN snails were used. Snails were infected as described above, using either SmBRE or SmVEN parasites. For each experimental infection, snails were recovered at 24 and 96 h after infection, and designated as follows: BB24 and BB96 for BgBRE infected with SmBRE; BV24 and BV96 for BgBRE infected with SmVEN; VV24 and VV96 for BgVEN infected with SmVEN; and VB24 and VB96 for BgVEN infected with SmBRE. Snails of each strain, BgBRE and BgVEN, were recovered and used as controls.

#### Hemocyte proliferation analysis: microscopy

Microscopic inspection of hemocyte proliferation was conducted using 12 infected BgBRE (6 BgBRExSmBRE and 6 BgBRExSmVEN) and 3 uninfected BgBRE snails. The hemocytes of 3 snails (biological replicates) were counted for each condition at 24h and 96h after infection. The proliferation of circulating hemocytes was studied by using a Click-iT EdU Alexa Fluor 488 Flow Imaging Kit (Molecular Probes). At each time point, circulating hemocytes were recovered by direct puncture after foot retraction and 1mM of EdU solution was added to the hemolymph. Three hours later, the amount of EdU incorporated by the circulating hemocytes was visualized *in-vitro* after fixation of the cells and performing a covalent coupling of Alexa Fluor 488 to the EdU residues trough a click chemistry reaction flowing manufacturer indications, then nuclei of hemocytes were counterstained with DAPI (Biotum) staining, and the sample was analysed on a confocal microscope using a Zeiss LSM 700, with 4 lasers (405, 488, 555 and 633 nm). Positive cells were counted and between-sample differences in the percentage of proliferation were tested using a Fisher exact test, with significance accepted at p-value<0.05.

#### Hemocyte proliferation and population profiles analysis: flow cytometry

Qualitative changes in hemocyte populations following infection by sympatric or allopatric parasites were studied using a flow cytometry approach. For this 72 infected BgBRE or BgVEN (36 infected by SmBRE and 36 infected by SmVEN) and 18 uninfected BgBRE or BgVEN snails were used. Six biological replicates (pools of 3 snails per replicate) were used for each condition. Flow cytometry was used to profile and assess the proliferation of circulating hemocytes using Click-iTEdUAlexa Fluor 647 labelling (Molecular Probes). At each time point, 1mM of EdU solution was injected into pericardial cavity of each snail. Three hours later six replicates of 3 snails were collected, and the hemolymph was extracted from the head-foot according to standard procedures [80]. The hemolymph was pooled from the three snails, and 100 µl were subjected to analysis with the above-listed kit, according to the manufacturer’s instructions. The percentage of proliferative cells was calculated by flow cytometry. The hemocytes were profiled along the course of infection using Side Scatter Chanel (SSC) to estimate cell granularity and Forward Scatter Chanel (FSC) to estimate cell size. The cell repartition along these two parameters enables to identify cell sub-populations. The flow cytometry was performed using a FACS Canto from BD Biosciences (RIO Imaging Platform, Montpellier, France). For each sample, 10,000 events were counted. The results were analyzed with the FlowJo V 10.0.8 software. Between-group differences in the percent of proliferation were tested using the Mann-Whitney U-test, with significance accepted at p-value<0.05.

### Histological procedures

A histological approach was conducted in order to investigate differences in miracidia to sporocyst development, while comparing sympatric and allopatric parasite growth, development and maturation into snail tissues. BgBRE snails were infected as described above with either 10 mi of SmBRE (sympatric) (n = 6 snails) or 10 mi of SmVEN (allopatric) parasite (n = 6 snails). At each time point, 24 and 96 h after infection, three snails were fixed in Halmi’s fixative (4.5% mercuric chloride, 0.5% sodium chloride, 2% trichloroacetic acid, 20% formol, 4% acetic acid and 10% picric acid-saturated aqueous solution). Embedding in paraffin and transverse histological sections (3-μm) were performed using the RHEM platform (Montpellier, France) facilities. The slides were stained using Heidenhain’s azan trichromatic staining solution as follows: (i) serial re-hydration was performed in toluene followed by 95%, 70%, and 30% ethanol and then distilled water; (ii) coloration was performed using azocarmine G (70% ethanol, 1% aniline, 1% acetic alcohol, distilled water, 5% phosphotungstic acid, distilled water, Heidenhain’s azan) and (iii) serial dehydration was performed using 95% ethanol, absolute ethanol, and toluene. The preparations were then mounted with Entellan (Sigma Life Science, St. Louis Missouri, USA) and subjected to microscopic examination. When a parasite is observed in snail tissue, the parasite size was measured using the imaging analysis software ImageJ (v2.0.0) for each adjacent histological section in which the parasite is observed. The contour of the parasite is detailed very precisely using ImageJ and the pixel number is reported on a size scale analyzed in the same manner to calculate parasite size. Size is expressed as parasite surface in μm^2^ as the mean of the 3 bigger parasite sections recorded. At 24h, n = 9 sympatric and n = 8 allopatric parasites were measured and at 96h, n = 7 sympatric and n = 10 allopatric parasites were measured. The size differences between sympatric and allopatric parasite groups were tested using the Mann-Whitney U-test with statistical significance accepted at a p-value < 0.05.

### In-silico characterization of Schistosoma miRNAs

Parasites may communicate or interfere with their host using different strategies based mainly on excreted/secreted products released into hemolymph. In this context, miRNAs appeared as the most relevant mean of communication that can be used by parasites. To test for such hypothesis *S. mansoni* miRNAs were analyzed *in-silico* by comparing the relevant miRNA database (miRBase) to our RNAseq libraries generated at the 24h following sympatric or allopatric infections. S. *mansoni* precursor sequences were downloaded from miRBase (http://www.mirbase.org, 03/09/2017), and high-quality reads from naive (BgBRE) and 24 h post-infection samples (BB24, BV24, BR24) were aligned against a *S. mansoni* sub-database of miRBase, as previously described [81]. The identified precursors were confirmed by alignment of high-scoring reads onto precursor and mature miRNAs from miRBase.

Solely reads with 100% identity were retained for analysis. The localization of each read against miRNA sequence allowed us to identify either the precursor or just the mature miRNA. Precursors found under both naive and infected conditions were excluded to retain exclusively the miRNAs present in samples from infected snails and avoid cross-species contamination because of the potential conserved features of miRNAs from *B. glabrata* and *S. mansoni*.

Putative miRNA targets were predicted from among the differentially represented immune-related transcripts (figure 1) using Miranda tools (using parameters: Miranda input_miRinput_Transcriptome - out results.txt -quiet -sc 140 -en −15) [82]. Because mature miRNAs may exist in two forms depending on which strand (5’-3’) of the precursor stem-loop is maturated the predicted interactions could involve the 5’ and/or 3’ forms, as noted. The results were extracted using the awk tool, listed in S4 Table, and used to generate a Venn diagram. To confirm the ability of a selected pre-miRNA to produce the stem-loop necessary to produce the mature form, the secondary structures of precursor were predicted using RNA structure Web tool (http://rna.urmc.rochester.edu/RNAstructureWeb, 03/09/2017) using default parameters.

### Snail survival analysis

Allopatric or sympatric parasites could have different levels of virulence or impacts on their host that could impair snail survival. To test for such discrepancy we investigated the mortality rates of infected snails over the course of sympatric or allopatric infections. One hundred and sixty BgBRE snails were infected as described above with SmBRE or SmVEN strains (n=50), and 60 non-infected BgBRE snails were retained as controls. The numbers of dead snails were compiled weekly for 14 weeks. A Kaplan-Meier estimator was used to estimate the survival function from lifetime data. Survival curves were generated using the xlstats Mac software and the log-rank test was applied with significance accepted at p<0.05.

## Acknowledgements

We thank Ms. Cécile Saint-Béat, Ms. Nathalie Arancibia, and Ms. Anne Rognon for their work and diligence in helping generate some of the data described herein.

## Supporting Information Legends

S1 Fig: Clustering of all differential represented transcripts

Clustering of differentially represented transcripts. Heatmap representing the profiles of the 1,895 differentially represented immune-related transcripts in the BB, BV, or BR interactions along the time course of infection (at 24 and 96 h). Each transcript is represented once and each line represents one transcript. Colors: yellow, over-represented transcripts; purple, under-represented transcripts; and black, unchanged relative to levels in control naïve snails.

S2 Fig: Mortality of B. glabrata snails after infections

The survival rates of *B. glabrata* subjected to infection by different *S. mansoni* strains were observed over 14 weeks. Kaplan Meier graphs were generated using xlstat, and the log-rank test (p < 0.05) was used to test for significant between-group differences. Colored curves indicate the mortality rates of naïve snails (yellow) (n=60), snails infected by the sympatric parasite (BB, BgBRE/SmBRE, blue) (n=50), and snails infected by the allopatric parasite (BV, BgBRE/SmVEN, green) (n=50). The difference in mortality between naïve and infected snails was significant (p<0.05), whereas that between the two infected conditions was not (p = 0.243).

S3 Fig: Blast-like cells are non-proliferative cells

*In vitro* EdU labeling of hemocytes collected for *in vitro* analysis. Confocal microscopy of EdU-labeled hemocytes from snails subjected to the allopatric interaction BgBRE/SmVEN at 24 h post-infection (BV24). Pictures corresponded to the merge of DAPI labelling (blue); EdU labelling (green) and phase contrast pictures. The white arrows indicate the Blast-like cells. Blast-like cells were never labelled by EdU, indicating that these cells are not proliferative when circulating in the hemolymphe. Three individual snails were used for each condition. Green label: EdU-positive cells; and blue label: DAPI. Magnification x63.

S1 Table: *Biomphalaria* and *Schistosoma* compatibility between African and South-American strains

The prevalence (P %: percentage of snail infected) and intensity (I: number of parasites per infected host) of infection are presented for each experimental infection. The indicated values correspond to 10 miracidia. Each pairwise combination of *Biomphalaria glabrata* (BgBRE, BgVEN), *Biomphalaria pfeifferi* from Senegal (BpSEN) and *Schistosoma mansoni* (SmBRE, SmVEN, SmSEN) or *Schistosoma rodhaini* (Srod) were tested for compatibility. The observation of cercariae shedding is also indicated. Cercariae shedding have been observed between 35 and 38 days after miracidial infections *NA*: non-available data.

S2 Table: List of differentially represented transcripts in RNAseq clusters.

Quality reads (Phred score >29) were aligned on the transcriptome assembly using the C++ script Bowtie2 (v2.0.2) (255 score) running thanks local engine using Galaxy Project server (Giardine, Riemer et al. 2005). The DESeq2 software (Love, Huber et al. 2014) (v2.12;http://www.bioconductor.org/packages/release/bioc/html/DESeq2.html) (defaults settings) allows for quantifying the differential gene expression with comparing two biological duplicates from uninfected snails sample (Bre1 and Bre2) against infected samples (Pvalue<0.1). For each cluster transcript ID, Blast2GO annotation and Log2FC results were indicated.

S3 Table: List of transcripts express by Schistosoma within Biomphalaria glabrata tissues highlight in RNAseq clusters.

The Biomphalaria (v1) and Schistosoma (v5.2) genome have been concatenate to mapped the RNAseq reads of each experimental condition. Only quality reads (Phred score >29) were aligned to the concatenate genomes using Bowtie2 (v2.0.2), which run locally on the Galaxy project server (Giardine, Riemer et al. 2005). The reads that mapped only once are conserved. Elimination of reads which mapped on Biomphalaria genome and only the reads that mapped on Schistosoma genome are kept.

S4 Table: miRNAs precursor identified in *Biomphalaria glabrata* RNAseq data.

